# Delta-band neural envelope tracking predicts speech intelligibility in noise in preschoolers

**DOI:** 10.1101/2023.02.22.529509

**Authors:** Tilde Van Hirtum, Ben Somers, Eline Verschueren, Benjamin Dieudonné, Tom Francart

**Author notes:** Correspondence: Tilde Van Hirtum, Herestraat 49, bus 721, 3000 Leuven, Belgium.

## Abstract

Behavioral tests are currently the gold standard in measuring speech intelligibility. However, these tests can be difficult to administer in young children due to factors such as motivation, linguistic knowledge and cognitive skills. It has been shown that measures of neural envelope tracking can be used to predict speech intelligibility and overcome these issues. However, its potential as an objective measure for speech intelligibility in noise remains to be investigated in preschool children. Here, we evaluated neural envelope tracking as a function of signal-to-noise ratio (SNR) in 14 5-year-old children. We examined EEG responses to natural, continuous speech presented at different SNRs ranging from -8 (very difficult) to 8 dB SNR (very easy). As expected delta band (0.5-4 Hz) tracking increased with increasing stimulus SNR. However, this increase was not strictly monotonic as neural tracking reached a plateau between 0 and 4 dB SNR, similarly to the behavioral speech intelligibility outcomes. These findings indicate that neural tracking in the delta band remains stable, as long as the acoustical degradation of the speech signal does not reflect significant changes in speech intelligibility. Theta band tracking (4-8 Hz), on the other hand, was found to be drastically reduced and more easily affected by noise in children, making it less reliable as a measure of speech intelligibility. By contrast, neural envelope tracking in the delta band was directly associated with behavioral measures of speech intelligibility. This suggests that neural envelope tracking in the delta band is a valuable tool for evaluating speech-in-noise intelligibility in preschoolers, highlighting its potential as an objective measure of speech in difficult-to-test populations.

## 1 INTRODUCTION

Globally, 34 million children (~0.2% of all children aged 0 to 18 years) have disabling hearing loss, i.e., hearing loss greater than 30 decibels (dB) in the better ear (report of World Health Organization, 2021). The prevalence rises to 5% when mild and unilateral hearing losses are also considered (Wang et al., 2019). Unaddressed hearing loss has been proven to affect children’s speech and language development, educational attainments and social skills. Through early detection and interventions many of these impacts can be mitigated, highlighting the importance of accurate hearing diagnostics (O’Donoghue, 2013). Evaluation of speech intelligibility is a fundamental component of hearing loss assessment and rehabilitation. It can determine speech intelligibility and discrimination of speech features, and provides insight into the perceptual abilities of an individual (Eggermont, 2017). The current gold standard in measuring speech intelligibility relies heavily on behavioral tests. While these tests are reliable and fast in healthy adults, it can be difficult to assess speech intelligibility in children. A child’s abilities and limitations in the language domain as well as their cognitive abilities strongly affect the results of a behavioral test. Moreover, the type of response task, the speech material, and children’s motivation and involvement should also be considered (Mendel, 2008). In addition, measures of speech intelligibility are often restricted to testing in quiet, due to a lack of appropriate and reliable speech in noise tasks (van Wieringen & Wouters, 2022). However, the high prevalence of noise in children’s natural listening environments, especially in settings where children learn and play (e.g., clamorous classrooms and rowdy playgrounds), poses serious challenges for a still immature auditory system (e.g., Ambrose et al., 2014; Neuman et al., 2010). Therefore it is essential that we can measure the ability to understand speech in these noisy, everyday environments.

An objective approach, using EEG to measure neural tracking in response to natural running speech, could overcome the current challenges in pediatric hearing assessment and provide a more reliable measure of a child’s speech intelligibility (for a review see Gillis et al., 2022b). Neural tracking refers to the process by which neural responses in a listener’s brain time-lock to dynamic patterns of the presented speech, such as the speech envelope. The speech envelope contains acoustical information (Rosen, 1992) and reflects phoneme, syllable and word boundaries (Peelle et al., 2013), which are critical for speech intelligibility (Shannon et al., 1995).

Indeed, a rich literature has found that during speech perception, auditory neural activity tracks the temporal fluctuations of the speech envelope in frequency bands matching the important occurrences of speech information, i.e., phrases/sentences (below 2 Hz) and syllables (2-8 Hz) (Vander Ghinst et al., 2019; Ding et al., 2016; Bourguignon et al., 2013; Ahissar et al., 2001; Gross et al., 2013; Meyer et al., 2017; Molinaro & Lizarazu, 2017). More importantly regarding diagnostic purposes, many researchers have shown that neural envelope tracking is affected by the intelligibility of the presented speech (Peelle et al., 2013; Gross et al., 2013; Di Liberto et al., 2018) and is significantly correlated with behavioral measures of speech intelligibility (Ding et al., 2014; Vanthornhout et al., 2018; Lesenfants et al., 2019a; Verschueren et al., 2021).

Most of the above-mentioned studies have been conducted in adults. Research on neural envelope tracking in children is scarce, despite the functional relevance that neural tracking might have for objectively measuring children’s speech intelligibility. Moreover, cross-sectional evidence shows that auditory neural activity changes drastically during childhood (e.g., Cragg et al., 2011; Vander Ghinst et al., 2019; Schneider & Maguire, 2019; Panda et al., 2020), which makes it difficult to extrapolate outcomes from neural tracking research in adults to children. Furthermore, previous research in children mostly involved older children (> 10 years) with dyslexia, focusing on the relation between neural tracking and reading development/experience. These studies show consistent coherence in the delta frequency range (0.5-4 Hz) (Molinaro et al., 2016) and theta (4-8 Hz) (Abrams et al., 2009; Molinaro et al., 2016) using natural speech. In addition, they found that dyslexic children have impaired tracking at low frequencies (< 2 Hz) compared to typically developing children (Molinaro et al., 2016; Power et al., 2013, 2016; Di Liberto et al., 2018; Destoky et al., 2022).

Although evidence of the neural tracking mechanisms in typically developing children is limited, more recent studies have been able to show successful neural tracking of continuous natural speech in both infants (< 1 year) (Kalashnikova et al., 2018; Tan et al., 2022; Attaheri et al., 2022) and young children between 4 and 9 years old (Vander Ghinst et al., 2019; Ríos-López et al., 2020; Tan et al., 2022). For example, Ríos-López et al. (2020) showed that speech-brain coupling already occurs at 4 years of age in the delta-band frequency range, but not at theta frequencies. Similarly, Vander Ghinst et al. (2019) found significant speech tracking in children aged 6-9 at < 1 Hz frequencies, while neural envelope tracking was reduced or even absent in the theta band.

Furthermore, neural envelope tracking in children has typically been evaluated under optimal, noiseless listening conditions. Findings from multiple, behavioral studies provided evidence that mature performance on a wide range of speech-in-noise measures is established by about 9-10 years of age, while younger children require a higher signal-to-noise ratio (SNR) to achieve adult-like performance (for a review see Leibold 2017). Thus, children seem more susceptible to the detrimental effects of noise on speech intelligibility. Only one study to our knowledge has studied neural tracking of speech-in-noise in typically developing children using magnetoencephalography (MEG). In accordance to behavioral measures, Vander Ghinst et al. (2019) found that neural tracking differs between typically developing children and adults and, that noise differentially corrupts neural tracking in children. In adults, their results showed a clear effect of noise at delta frequencies (<1 Hz), that is, a decrease in coherence as SNR decreased and speech is less intelligible. However, in children increasing noise decreased coherence more strongly than in adults. Additionally, children’s coherence was drastically reduced or even absent in comparison with adults in the theta band regardless of SNR. Generally, these results are in line with previous behavioral studies showing children’s poorer speech intelligibility in adverse listening conditions (Johnson, 2000; Talarico et al., 2007; Neuman et al., 2010; Leibold, 2017). However, we cannot conclude from this study that delta and/or theta coherence is directly related to speech-in-noise intelligibility, since only indirect behavioral measures (e.g., intelligibility rating) were included. In addition, MEG-based recordings were used and various practical aspects of MEG in its current form (e.g., cost) pose a limitation to its large-scale usability in clinical practice, compared to electroencephalography (EEG) (Destoky et al., 2019).

Our recent research provides a framework for investigating neural tracking of different speech features, including the speech envelope (Lesenfants et al., 2019b; Verschueren et al., 2021; Gillis et al., 2022b), using EEG. The approach combines both linear decoding (backward-modelling) and encoding (forward-modelling) models, providing complementary information about neural tracking. The backward model is a model to reconstruct the speech envelope from the associated EEG recording, whereas the forward model predicts the EEG responses to speech and can be used to study the spatio-temporal dynamics of the response similarly to event related potentials (ERPs). This is the first study to use this method in preschoolers and we aim to investigate the validity of a measure that has previously been used only with adults (e.g., Vanthornhout et al., 2018). More specifically, in this study, we investigate (i) the effect of SNR on neural envelope tracking in preschoolers, and (ii) whether neural envelope tracking reflects speech intelligibility by evaluating the correspondence between neural envelope tracking and behavioral measures of speech intelligibility in noise. We have two specific hypotheses. First, we predicted that as SNR increases, neural envelope tracking increases, given that previous research has shown that stronger neural responses are associated with better speech intelligibility (Ahissar et al., 2001; Peelle et al., 2013; Ding et al., 2014; Vanthornhout et al., 2018). Secondly, we hypothesize that, similar to studies in adults (Vanthornhout et al., 2018; Lesenfants et al., 2019a), behaviorally measured speech intelligibility in noise is significantly correlated with our neural, objective measure.

## 2 MATERIALS AND METHODS

### 2.1 Participants

Fourteen 5-year-old children, recruited from the third year of kindergarten, participated in the experiment (6 female). All children were native Dutch speakers, had normal peripheral hearing (hearing thresholds ≤ 30 dB HL for frequencies from 0.5 to 4 kHz) and had normal or corrected to normal vision. None of the children were at risk for a cognitive or language delay nor had a family history of developmental disorders, as reported by the parents. The study protocol was approved by the local Medical Ethics Committee (reference no. S57102) and all parents provided written informed consent before the experiment. All children received a gift voucher for participating.

### 2.2 Behavioral measurements

#### 2.2.1 Speech-in-noise intelligibility

Speech intelligibility in noise was assessed using the Leuven Intelligibility Peuter Test (Lilliput; van Wieringen & Wouters, 2022). This test consist of 20 lists of 11 three-phoneme consonant-vowel-consonant words (e.g., “bus”) uttered by a female speaker. All lists were presented in a stationary speech-weighted noise, matching the long-term average spectrum of the speaker. The noise level was fixed at 65 dB SPL, whereas the speech level was adjusted to obtain the targeted SNRs. Each child started with 1 training list at 0 dB SNR. Thereafter, different lists were presented monaurally to the right ear at four fixed SNRs: 0, −3, −6 and −9 dB SNR. If necessary, additional lists at −12 dB SNR (i.e., if scores at −9 dB SNR >50%) were presented. Two lists were presented per SNR. Children were instructed to recall each word as accurately as possible. The result of the Lilliput test is a phoneme score, i.e. the number of phonemes correctly repeated by the child. The average phoneme score per SNR was used for further analysis. Words were played through Peltor H7A headphones using the software platform APEX (Francart et al., 2008) on a Samsung Galaxy Tab A tablet.

#### 2.2.2 Receptive language

Children’s receptive language skills were examined using the Peabody Picture Vocabulary Test-III-NL (PPVT-III-NL; Dunn & Dunn, 1997). The PPVT-III-NL was administered in order to confirm the absence of a language impairment. It is a norm-referenced test that consists of selecting 1 of 4 pictures corresponding to a given word, at increasing levels of difficulty. Raw scores were calculated as the total responses correct and converted to age-adjusted standard scores (100 ± 15; mean ± SD) according to the standard values included in the test manual. The mean of the PPVT-III scores was 114 (SD = 12.82), ranging from 86 to 127 showing that all children performed well within age norms.

### 2.3 EEG experimental procedure

#### 2.3.1 Speech material and procedure

During the EEG recordings, children listened to four different stories of “Little Polar Bear”, the children’s series by Hans de Beer, narrated by the same native Flemish speaker as the Lilliput. Each story was 10 to 12 minutes long and segmented in two-minute fragments taking sentence boundaries into account. The stories were presented in a stationary speech-weighted noise, obtained by the long-term average spectrum of all stories together. For all stories the speech level was fixed at 60 dB A, while the noise level was adjusted to obtain five SNR conditions: 8, 4, 0, −4 and −8 dB SNR. Every two-minute segment was presented randomly at a different SNR. In total, every SNR was presented three times (i.e., 6 minutes), with the exception of the easiest SNR condition (8 dB SNR). The 8 dB SNR condition was presented for an additional 6 minutes at the end of the recording session.

#### 2.3.2 Data acquisition

EEG signals were recorded using a Biosemi ActiveTwo System at a sample rate of 8192 Hz. This system uses 64 Ag/AgCl electrodes, distributed over the scalp according to the international 10-20 system. The electrode offsets were kept between −30mV and 30mV to ensure stable recording.

All recordings were carried out in a soundproof and electrically shielded room, using a child-friendly and age-appropriate protocol. Children were comfortably seated in a chair approximately 1 m from a sound-permeable screen. Speech stimuli were presented at a sample rate of 48000 Hz through a GENELEC (8020A) loudspeaker positioned at head-height of the seated child using the software platform APEX (Francart et al., 2008) and an RME Fireface UC soundcard (Haimhausen, Germany). To mimic a storytelling scenario children watched images of the corresponding “Little Polar Bear” books projected on the screen in front of them while listening to the stories.

Children were instructed to attend to the stories and to avoid moving as much as possible. Additionally, children were accompanied by an experienced test leader in the EEG cabin to monitor alertness and movement. During each story, each child was asked 3 multiple-choice comprehension questions to improve attention. Short recesses between each two-minute segment and each different story were also used to motivate the children and, if necessary, allow them to rest or move before resuming the story.

### 2.4 EEG data analysis

We measured neural tracking of the speech envelope as a function of stimulus SNR using an envelope reconstruction approach (backward-modelling) and temporal response function estimation (forward-modelling). All signal processing was performed offline using Matlab R2016b (The MathWorks Inc, 2016).

#### 2.4.1 Envelope reconstruction

To measure neural envelope tracking, we used the envelope reconstruction approach described in detail by Vanthornhout et al. (2018) and Verschueren et al. (2020). In summary, the speech envelope was extracted according to Biesmans et al. (2017) who showed good reconstruction accuracy using a gammatone filter bank followed by a power law. The acoustic envelope was then downsampled from 48000 to 256 Hz in order to decrease computation time. Next, the speech envelope was band-pass filtered between 0.5-4 Hz (delta band) and 4-8 Hz (theta band) using a Chebyshev filter (with 80 dB attenuation and 10% outside the passband) and further downsampled to 128 Hz. Similarly to the speech envelope, the EEG data was first downsampled from 8192 Hz to 256 Hz. Next, a multi-channel Wiener filter (Somers et al., 2018) was applied to the EEG data to remove common EEG artifacts such as eye blinks and muscle artifacts. Bad EEG channels were interpolated from their 5 neighbouring channels using Fieldtrip (Oostenveld et al., 2011). We then re-referenced each EEG signal to a common-average reference. Finally, the EEG data was band-pass filtered similarly to the speech envelope and further downsampled to 128 Hz.

For each SNR condition, the reconstructed envelope is obtained by applying a condition-specific linear decoder, calculated using ridge regression as implemented in the mTRF toolbox (Lalor et al., 2006). Each decoder is basically a spatiotemporal filter that combines the 64 EEG channels and their time-shifted versions from 0 to 250 ms (the integration window) into a reconstruction of the envelope. After normalization, envelope decoders were trained using a leave-one-out 6-fold cross-validation scheme: for each SNR condition, containing 6 minutes of EEG data, 5 minutes were used to train the decoder, which was then applied on the EEG of the remaining minute to obtain a 1-minute envelope reconstruction. This was repeated 6 times to obtain envelope reconstructions for all folds. All reconstructions within a SNR condition were concatenated and compared to the original envelope using a bootstrapped Spearman correlation, resulting in an envelope tracking value for that SNR condition. The significance level of this correlation value is calculated by constructing a null distribution by correlating 1000 random permutations of the real and reconstructed envelopes with each other, and taking the 2.5 and 97.5 percentiles to obtain a 95% confidence interval.

#### 2.4.2 Temporal response function (TRF) estimation

The envelope reconstruction approach is a powerful analysis tool, which integrates information of multiple EEG channels and their time-shifted versions to reconstruct the envelope. However, this type of analysis does not allow an interpretation of the spatial pattern of the response (Haufe et al., 2014). We therefore conducted a linear forward modelling approach, in addition to the envelope reconstruction approach. This forward modelling approach predicts EEG given a speech representation and results in a temporal response function (TRF) for each channel. A TRF represents an impulse response function of how the brain responds to the stimulus envelope. The main advantage of estimating TRFs is that their morphology provides valuable insights concerning the neural response latency, amplitude and topology across the scalp. The first signal processing steps are identical to the envelope reconstruction approach. Only the band-pass filtering was carried out within a broader frequency band (0.5-25 Hz), after which the EEG data is normalized. As with the backward-modelling approach, TRFs are computed using 6-fold cross-validation for every SNR condition, where 5 minutes are used for training and 1 minute for validation. TRFs are computed for every channel using the boosting algorithm as described by David et al. (2007), implemented in the python Eelbrain toolbox (Brodbeck, 2020). In short, this TRF estimation algorithm works by starting from an all-zero TRF model and iteratively improving this model by adding a weight at a latency that causes the largest improvement in the response prediction. The iterations terminate when no significant improvements to the model can be made anymore for a given step size by which the weights are changed. The advantage of this method is that the resulting TRF is sparse, i.e., for a well-chosen step size, only the peaks that contribute most to the response prediction are included in the TRF while other TRF weights are zero. TRFs were computed over a latency range (integration window) between −200 and 500 ms with an adaptively decreasing stepsize. For further analysis and visualisation, the estimated TRFs were convolved with a Gaussian kernel of 9 samples long (SD=2) to smooth over the time lags. Next, smoothed TRFs were averaged across all folds.

#### 2.4.3 Reliability of envelope reconstruction scores

To investigate potential changes in neural tracking over the course of the recording session, we calculated a measure of reliability. In particular, test-retest reliability can be used to reflect the variation in multiple measurements on the same subject under the same conditions. Here we quantified test-rest reliability by means of intraclass correlation (ICC) (Shrout PE, 1979), which is one of the most commonly used reliability measures in the neuroimaging field (Bennett & Miller, 2010). In this study, every SNR was presented three times (i.e., 6 minutes), with the exception of the easiest SNR condition. For the 8 dB SNR condition, an additional 6 minutes were presented at the end of each measurement. Test-retest reliability was estimated between the envelope reconstructions scores for the first and last 6 minutes, under the assumption of a single-measurement, absolute-agreement, two-way mixed model, i.e., ICC(A,1) (Koo & Li, 2016). ICC estimates and their 95% confidence intervals were calculated using the irr package (Gamer et al., 2019).

### 2.5 Statistical analysis

Statistical analysis were carried out in R (version 4.0.3; R Core Team, 2020). All tests were performed with a significance level of *α* = 0.05 unless otherwise stated.

To investigate behavioral speech intelligibility in noise, the speech reception threshold (SRT; the SNR corresponding to a 50% percentage-correct score) and slope were determined. For each child individually a sigmoid function was fitted to their average phoneme scores using the following formula:

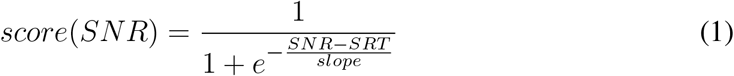

To assess the relation between stimulus SNR and neural envelope tracking, we fitted a linear mixed effect model (LME) per filterband using maximum likelihood criteria with the nlme package (Pinheiro et al., 2022). The following general formula was used:

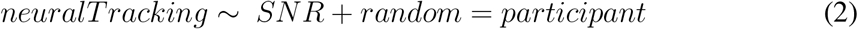

where “neuralTracking” represents the Spearman correlation between the real and the reconstructed envelope, TRF amplitude or latency depending on the outcome measure being investigated. In addition, a random intercept per participant was included to account for dependencies between measures from the same child. Residual plots of the selected model were analyzed to assess the assumption of normality and did not reveal any violations. All significant effects are discussed in the results section by reporting the *β* estimates, with the corresponding standard error (SE), degrees of freedom (df) and test statistics (*t*-value and *p*-value). Where applicable, post-hoc comparisons were carried out using a non-parametric Wilcoxon signed-rank test (2-tailed, *p* < 0.01), with Bonferroni correction for multiple comparisons.

Finally, the Spearman correlation between the envelope reconstruction scores and behaviorally measured speech intelligibility was computed to examine the relation between neural envelope tracking and speech intelligibility. Since different SNR conditions were used for the behavioral experiment, we estimated the percentage-correct scores corresponding to the SNR conditions (i.e. −8, −4, 0, 4 and 8 dB SNR) used for the EEG measurement from the individually fitted performance intensity functions.

## 3 RESULTS

### 3.1 Behavioral speech intelligibility

For each child, we fitted a psychometric curve through their average percentage correct scores per SNR from which the SRT and slope were derived. The mean of the individual behavioral SRTs was −5.52 dB SNR (SD = 0.94 dB), with an average slope of 7.99%/dB (SD = 2.39%/dB). Individual speech intelligibility scores evaluated at the different SNR conditions are shown in in Figure 1 together with the fitted performance intensity function on the group average and individual SRT estimates.

**Figure 1.**
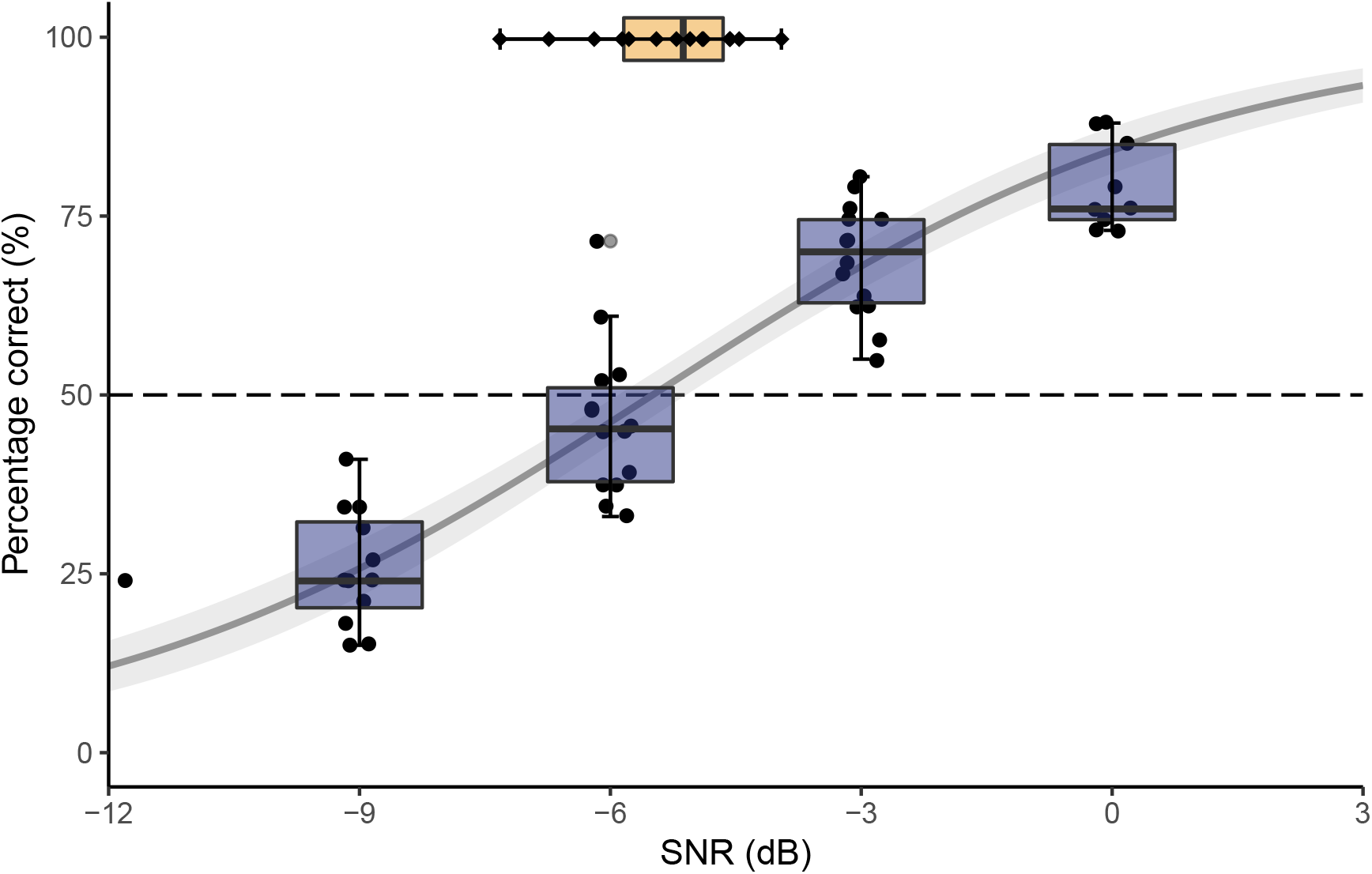
Behavioral speech intelligibility. Individual’s speech intelligibility (percentage correct) as a function of stimulus SNR and the fitted performance intensity function of the group average. Black dots represent the individual scores. The grey line shows the sigmoid fitted to the data ± the error of the fit of the model. The orange boxplot on top shows the distribution of the SRT. Individual SRT estimates are presented as black diamonds to show variation.

### 3.2 Envelope reconstruction

#### 3.2.1 Effect of SNR on envelope reconstruction

Figure 2 shows neural tracking of the speech envelope as a function of the SNR using a 0-250 ms integration window in the delta (0.5-4 Hz) and theta band (4-8 Hz). These results were obtained by a leave-one-out cross-validation approach within each 6-minute SNR block.

**Figure 2.**
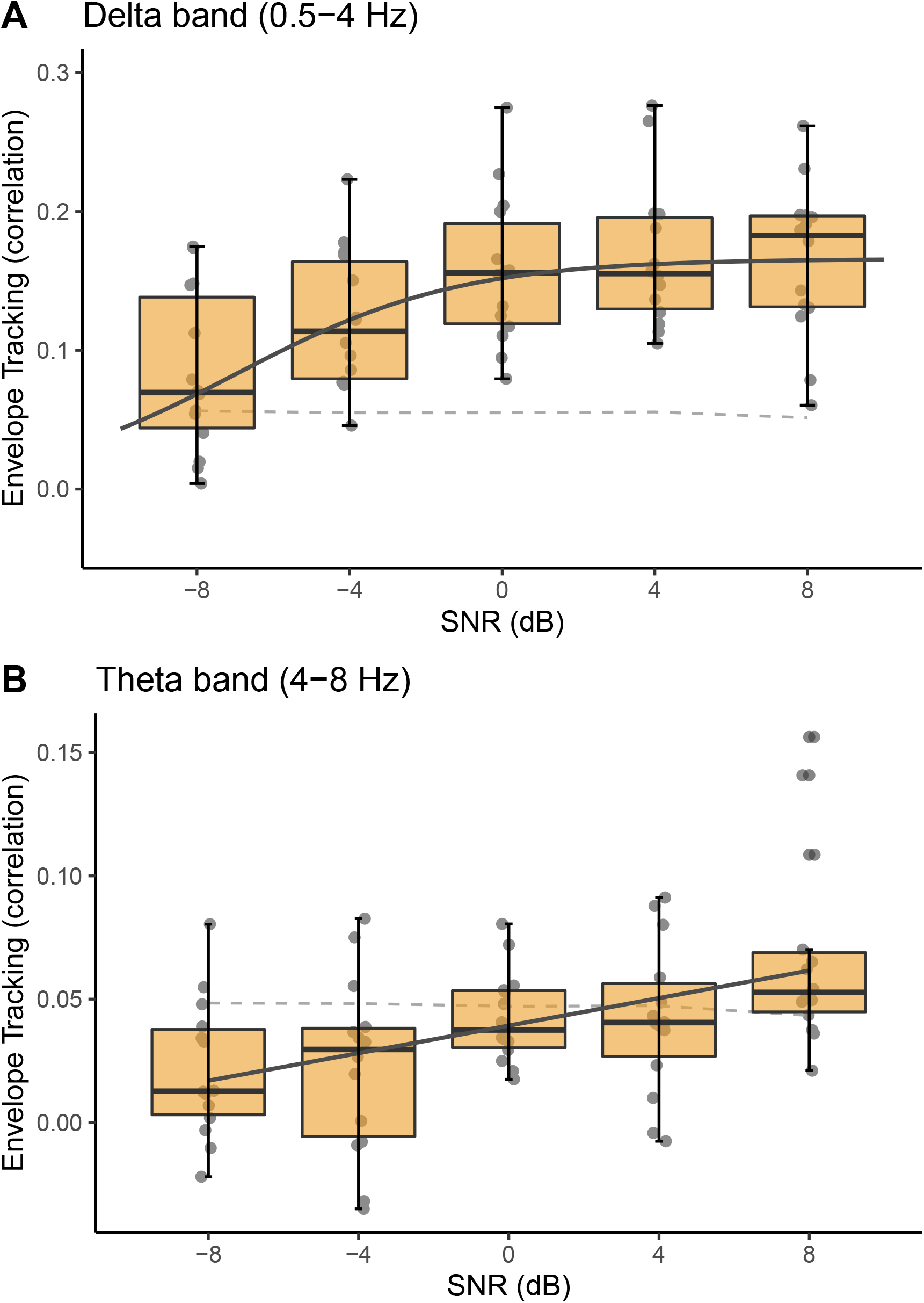
Neural envelope tracking as a function of stimulus SNR. **A** In the delta band, neural envelope tracking increases with increasing SNR until reaching a plateau around 0 dB SNR. The individual data points in each boxplot represent individual envelope reconstruction scores (i.e., correlation) to show variation. The solid line (dark grey) shows the sigmoid fitted to the data. The dashed line (light grey) shows the significance level of the envelope reconstruction scores. **B** In the theta band, neural envelope tracking monotonically increases with increasing SNR. However, limited statistically significant correlations were found below 8 dB SNR.

In the delta band, an increase of neural envelope tracking with increasing SNR was found (*b* = 0.005, SE = 0.0006, *p* < 0.001). However, we observed that neural tracking did not show a strictly monotonic increase with SNR. To investigate in more detail if neural tracking differed significantly between adjacent SNR conditions in the delta band, post-hoc comparisons were carried out using a non-parametric Wilcoxon signed-rank test (2-tailed, *p* < 0.01), with Bonferroni correction for multiple comparisons. Comparisons between increasing SNR conditions demonstrated that neural tracking increased significantly from −8 to −4 dB SNR (*p* = 0.003; *r* = −0.792) and from −4 to 0 dB SNR (*p* = 0.007; *r* = −0.724), but not in the two conditions with the lowest noise (0 to 4 dB SNR: *p* = 0.216, 4 to 8 dB SNR: *p* = 0.808). Therefore, a sigmoid function was fitted on the data across all subjects. Its midpoint was determined at −6.98 dB SNR and, convergence of the function (as calculated by the 95^th^%-value of the fit) was found around 1.5 dB SNR suggesting that neural tracking indeed increases with SNR until reaching a plateau between 0 and 4 dB SNR.

In the theta band, neural tracking again increased with SNR (*b* = 0.002, SE = 0.0006, *p* < 0.001). However, statistically significant responses were limited. Statistically significant neural tracking was observed at 8 dB SNR in 11 out of 14 children, at 4 and 0 dB SNR in 5 out of 14 children, and at −4 and −8 dB SNR in only 3 out of 14 children. Whereas in the delta band, statistically significant neural tracking was found in 14 of 14 children down to 0 dB SNR, at 13 of 14 children at −4 dB SNR, and at 9 of 14 children at −8 dB SNR. For further analysis, our frequency range of interest was therefore restricted to the delta band.

#### 3.2.2 Test-retest reliability of envelope reconstruction scores

Test-retest reliability of neural tracking over the course of the recording session was determined by means of ICC(A,1) for the first and last 6 minutes of speech presented at 8 dB SNR. ICC estimates were calculated based on a single-measure, absolute-agreement, 2-way mixed-effects model. We found a high degree of test-retest reliability of the envelope reconstruction values in the delta band. The average measure ICC was 0.876, with a 95% confidence interval from 0.629 to 0.96 (F(13,10.2) = 18.3, *p* < 0.001).

#### 3.2.3 Effect of speech intelligibility on envelope reconstruction

In order to assess the relation between speech intelligibility and neural envelope tracking, we calculated the percentage-correct scores for each SNR condition used in the EEG experiment using the individual psychometric curves. Figure 3 shows that neural tracking in the delta band increased with increasing speech intelligibility (*r* = 0.498; *p* < 0.001, Spearman rank correlation), suggesting that the better a child can understand speech, the higher the neural envelope tracking for that child.

**Figure 3.**
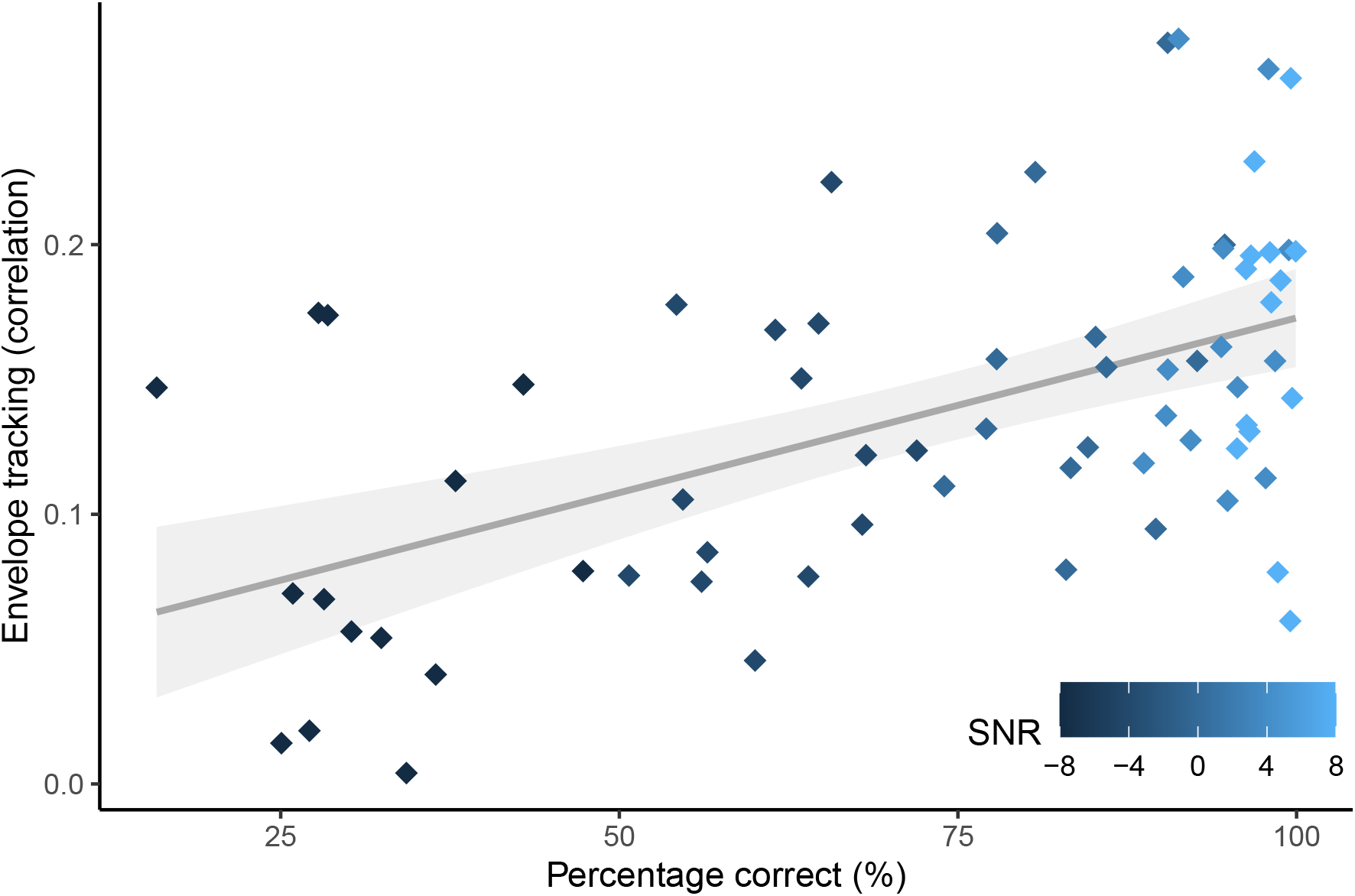
Relation between behavioral speech intelligibility and envelope tracking across all SNRs in the delta band. Envelope tracking increases with increasing speech intelligibility. The color gradient was used for illustrative purposes to mark SNR. The grey line shows the linear model fit. The shaded area represents the 95% confidence interval for the predictions of the linear model.

### 3.3 Effect of SNR on TRFs

The envelope reconstruction analysis showed an S-shape tendency of neural tracking as a function of SNR. That is, a monotonic increase in envelope tracking over SNR, until saturating between 0 and 4 dB SNR. To better understand the spatio-temporal properties of this effect, we calculated TRFs of every 6-minute segment (i.e., each SNR condition). Based on visual inspection of the topographies, 12 frontocentral channels were selected (presented by the black dots in Figure 4B) and averaged per subject, resulting in one TRF per SNR for each subject. Figure 4A shows a prominent peak appearing around ~83 ms.

**Figure 4.**
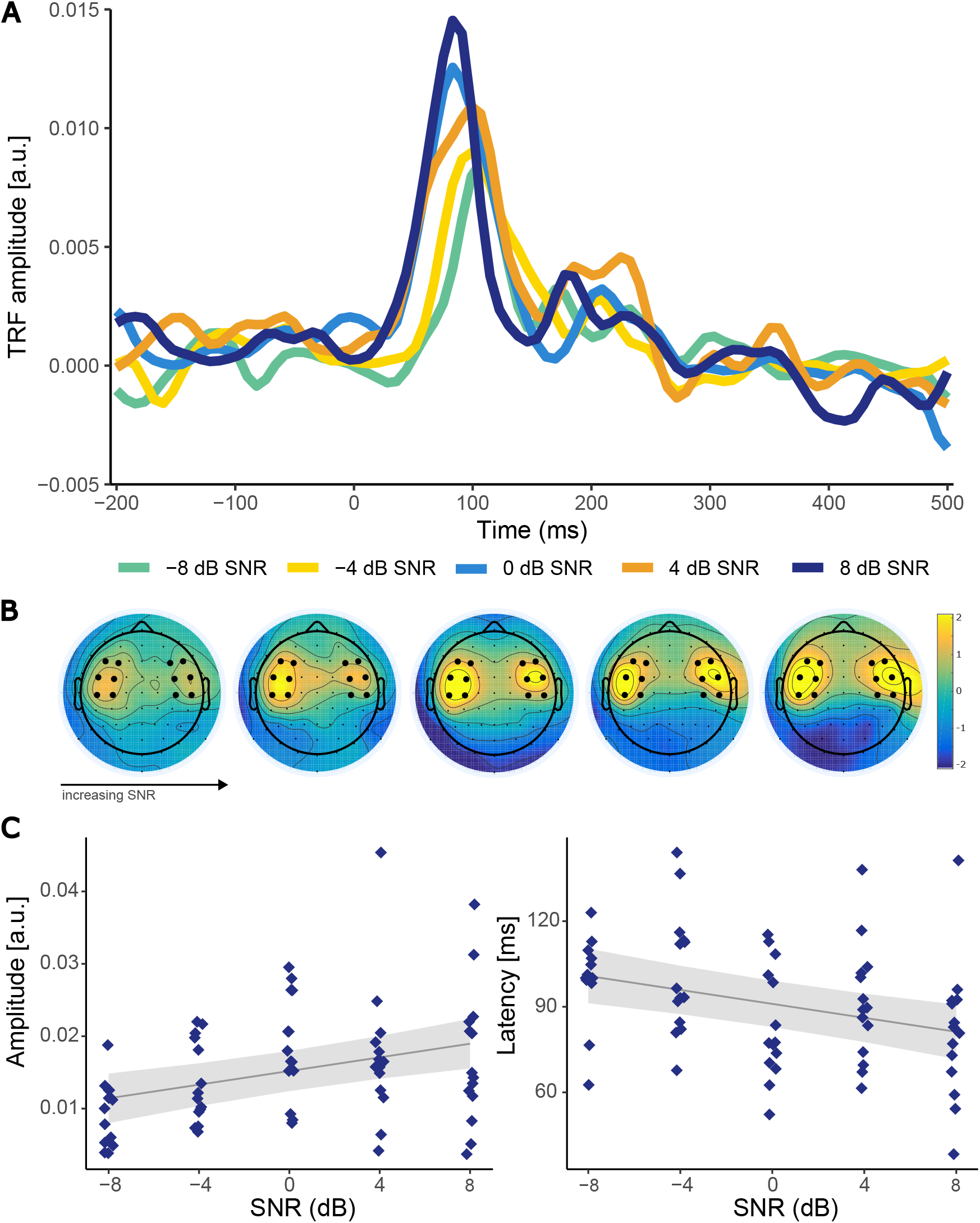
TRFs as a function of stimulus SNR. **A** Mean TRF activity over participants per SNR. **B** Topographies showing the associated peak topographies in the mean TRF (~70-90 ms) per SNR. Black dots represent the centro-frontal channels selected to calculate the TRFs in the time-domain in panel A. **C** The effect of SNR on peak amplitude (left) latency (right) of the TRF. Every diamond (blue) represents an individual participant. The grey line shows the linear model fit. The shaded area represents the 95% confidence interval for the predictions of the linear model.

The topography of this peak (70-90 ms) is shown in Figure 4B. To investigate the influence of stimulus SNR on the peak latency and amplitude in more detail, the maximum TRF amplitude for every subject per SNR was calculated between 0 and 150 ms. As shown in Figure 4C and confirmed by statistical analysis, with increasing SNR, the peak amplitude increases (*b* = 4.71 × 10^-4^, SE = 1.27 × 10^-4^, *p* < 0.001). In contrast, the latency of this peak decreases with increasing SNR (*b* = −1.22, SE = 0.32, *p* < 0.001). Post-hoc comparisons between the adjacent SNR conditions demonstrated that amplitude increased significantly from −4 to 0 dB SNR (*p* = 0.029; *r* = −0.582), whereas latency decreased from −4 to 0 dB SNR (*p* = 0.002; *r* = −0.808). We did not find significant differences between the two noisiest (−8 to −4 dB SNR: *p* > 0.05) nor the two conditions with the lowest noise (0 to 4 dB SNR: *p* > 0.05, 4 to 8 dB SNR: *p* > 0.05)

## 4 DISCUSSION

We examined the influence of stimulus SNR on neural envelope tracking of natural, continuous speech in young children. We recorded EEG in 14 normal-hearing pre-schoolers while they listened to age-appropriate stories at different stimulus SNRs. Neural tracking was investigated using both envelope reconstruction and TRF analysis. Our findings demonstrated that neural tracking of the speech envelope in the delta band (0.5-4 Hz) increased monotonically with increasing SNR, until stabilizing between 0 and 4 dB SNR, suggesting that neural envelope tracking remains stable as long as speech is intelligible. By contrast, neural tracking in the theta band (4-8 Hz) appeared to be more sensitive to noise. As SNR increased, neural tracking increased. However it is important to note that our results in the theta band did not fully indicate the presence of significant envelope reconstruction scores below 8 dB SNR. Lastly, we found that neural envelope tracking in the delta band was related to behavioral speech-in-noise performance, suggesting its potential for a realistic and objective measure of speech intelligibility in clinical practice.

We hypothesized that neural tracking of the speech envelope would increase as SNR, and thus speech intelligibility, increases. We demonstrated that this is indeed the case. However, in the delta band, we did not find a strictly monotonic increase of neural envelope tracking with SNR. As the SNR decreased, envelope reconstruction scores remained stable until <0 dB SNR, showing a S-shape correlation over SNRs (i.e., flat, followed by a decrease as SNR further decreases). Our finding aligns with previous results reported by Ding & Simon (2013) in which the effect of noise on neural entrainment to slow temporal modulations (<4 Hz) was investigated. Based on MEG recordings, they found that neural responses in the delta band remained stable as the SNR decreased, until the noise background is more than twice as strong as the speech. In accordance, a more recent MEG study including children aged 6-9 (Vander Ghinst et al., 2019) found no difference in coherence to speech in different levels of background noise (SNR ranging from +5 to −5 dB) at 1-4 Hz. Additionally, these results are consistent with the previous EEG work (Iotzov & Parra, 2019), including our own (Vanthornhout et al., 2018; Lesenfants et al., 2019a; Decruy et al., 2019), yielding an increase in neural tracking with SNR. However, Lesenfants et al. (2019a) reported a monotonic increase in the delta band, with neural tracking further increasing from - 1 dB SNR to noiseless conditions, thus only reaching a maximum in quiet. The discrepancy with our current results could be attributed to the differences in experimental paradigm. Previous studies (e.g., Vanthornhout et al., 2018) used a 15-minute-long recording of a story presented without noise to train a linear decoder. Then, this decoder was applied to shorter 2-minute-long segments presented at different SNRs. A recent study (Verschueren et al., 2021), however, demonstrated that reconstruction accuracy changes when using different stimulus intensities to train and test the decoder on. This suggests that a decoder is optimized to decode brain responses presented at the intensity it was trained on. Thus, when training and testing on the same stimulus parameters, neural tracking seems robust to stimulus intensity.

Taken together, we hypothesize that acoustically degrading the speech signal, such as by adding background noise, does not influence neural tracking provided that speech remains intelligible. It is well known that speech intelligibility and SNR are highly correlated. Therefore, it can be challenging to disentangle to what extent a difference in neural tracking arises from a change in the acoustics, such as different levels of background noise, or changes in speech intelligibility itself. However, our results revealed that changes in auditory stimulus characteristics (i.e., SNR) did not necessarily alter neural tracking, supporting this hypothesis. Similarly to our electrophysiological measures, behaviorally measured speech intelligibility starts to decrease significantly below 0 dB SNR, whereas average scores at 0 dB SNR exceeded 75%, reflecting good speech intelligibility for these young children (van Wieringen & Wouters, 2022). Thus, when acoustical degradation of the speech signal does not reflect a significant change in speech intelligibility, neural tracking in the delta band seems insensitive to stimulus SNR.

In contrast, neural envelope tracking in the theta band was influenced by stimulus SNR even though speech intelligibility remained high, replicating studies performed in adults (Ding & Simon, 2013; Verschueren et al., 2021). Furthermore, the number of children with significant tracking decreased with increasing noise level, suggesting that theta band tracking was more sensitive to SNR compared to the delta band. With limited noise present (i.e., 8 dB SNR), 11 of 14 children (delta band: 14 of 14 children) showed significant neural tracking of the speech envelope, whereas only 3 of 14 children (delta band: 9 of 14 children) still had significant neural responses when more noise was added (i.e., −4 dB SNR). This finding seems inconsistent with previous studies reporting significant theta tracking in adults (Bourguignon et al., 2013; Ding & Simon, 2013; Molinaro et al., 2016; Vanthornhout et al., 2018; Vander Ghinst et al., 2019; Lesenfants et al., 2019a). Yet, more recent results in young children also failed to demonstrate significant responses in the theta range (Vander Ghinst et al., 2019; Ríos-López et al., 2020).

The discrepancy between frequency bands could be rooted in developmental reasons such as the language acquisition stage of pre-school children. Previous behavioral results show that children’s sensitivity to phonological units progresses from larger to smaller units, thus slow to fast information in the speech signal, as linguistic and reading skills improve (Ziegler & Goswami, 2005). However, several studies found evidence of an early-developed ability to discriminate syllables already in preterm infants (Mahmoudzadeh et al., 2013; Nittrouer, 2006). Therefore, this hypothesis is at odds with the view that tracking at theta frequencies correspond to the processing of syllabic units (e.g. Ding & Simon, 2012b; Gross et al., 2013; Ding et al., 2016). Additionally, theta responses are generally lower compared to delta tracking (e.g. Vander Ghinst et al., 2019; Verschueren et al., 2021), supporting an alternative hypothesis that delta and theta tracking reflect different speech perception mechanisms. That is, delta band neural tracking involves more higher-order speech specific processing, whereas theta band activity is more sensitive to acoustic processing of the speech signal (Molinaro & Lizarazu, 2017; Ding et al., 2016)

This explanation can be further strengthened by the fact that we found a significant correlation between speech intelligibility and neural envelope tracking in the delta band. This is in line with previous studies (Meyer et al., 2017; Vanthornhout et al., 2018; Verschueren et al., 2021, 2019; Etard & Reichenbach, 2019) consistently reporting that increased neural envelope tracking reflects better speech intelligibility. This is especially promising for clinical applications in hearing loss assessment and rehabilitation.

In addition to the envelope reconstruction analysis, our TRF analysis revealed one prominent positive TRF peak around 70–90 ms. In contrast, the TRF in adults typically contains two distinct peaks (e.g., Ding & Simon, 2012a, 2013; Vanthornhout et al., 2019; Verschueren et al., 2021, 2022). The first peak, here called the P1, occurs relatively fast with latencies around ~50 ms and is thought to reflect mainly acoustic processing, whereas the second peak occurs at longer latencies (~ 100–150 ms; N1), and might be influenced by top–down processing related to speech intelligibility and attention (Brodbeck & Simon, 2020). In line with observations of cortical auditory evoked potentials (CAEP), we believe the positive peak in this study could be related to the P1 as reported in adults. In general, CAEP traces in young children can be characterized by a large positive peak (P1), emerging between 100–150 ms, whereas components N1 and P2 emerge more gradually with maturation and might not be reliably evoked until the age of 7–9 years old Albrecht et al. (2000); Ponton et al. (2000); Ceponien et al. (1998). Besides the change in temporal pattern of the CAEP (e.g., peaks emerging with increasing age), peak latency decreases with age (Wunderlich et al., 2006). Therefore, age–related changes reflecting maturation of auditory neural processing might explain why our results did not reveal a clear N1 or P2 component.

To further investigate how SNR affects the TRFs, we compared the individual peak amplitudes and latencies at the different stimulus SNRs. Our findings demonstrated that with increasing SNR, the amplitude of P1 increased. Furthermore, the latency of P1 decreased as the SNR increased. These results dovetail with previous research with adult participants showing that the amplitude of the early TRF peak (~50 ms) decreases, while the latency increases continuously with SNR (Ding & Simon, 2013; Jan et al., 2022). A delay in neural responses has been hypothesized to reflect a decrease in neural processing efficiency (Bidelman et al., 2019; Gillis et al., 2022a). In addition, increased latencies have previously been related to increasing task demand such as lower stimulus intensity, increasing background noise or vocoded speech both for neural processing of continuous speech (Mirkovic et al., 2019; Verschueren et al., 2021; Kraus et al., 2021) as well as simple sounds (Billings et al., 2015; Van Dun et al., 2016; Maamor & Billings, 2017; McClannahan et al., 2019). In the current study, we observed the same effect of background noise in pre-school children. However, the results in this study only showed significant differences in peak amplitude and/or latency from −4 to 0 dB SNR and not between the lower noise conditions, similar to the previously discussed envelope reconstruction results. This, again, suggests that decreasing the stimulus SNRs only influences the amplitude and latency of the neural response if speech intelligibility is affected. This aligns with the finding of Verschueren et al. (2021) that the P1 latency increased when audibility of the stimulus decreased, but only when audibility affected speech intelligibility and listening effort.

### 4.1 Limitations and future directions

It is important to note that, in this study particular emphasis has been placed on neural tracking of the speech envelope. Even though the envelope is essential for speech intelligibility (Shannon et al., 1995), only taking into account basic acoustic properties as a measure of speech intelligibility might be an oversimplification. As reviewed in detail by Brodbeck & Simon (2020), neural responses can be predicted more accurately when not only considering other acoustic features, but also higher-level linguistic speech representations. Therefore, neural envelope tracking should be considered a measure to assess whether the pre-conditions for speech intelligibility are met, rather than a direct measurement of speech intelligibility. Nevertheless, neural envelope tracking is shown to be quite robust, therefore remaining a good candidate for use in a clinical setting.

A second caveat of this study is that we included normal-hearing children as a first cohort, since speech intelligibility in noise had never been evaluated in such young children. However, for clinical applications of our measure, future research should study the impact of parameters such as hearing loss and hearing aid fitting. A number of studies have already demonstrated the use of CAEPs for validation of hearing aid fitting in hearing-impaired children (e.g., Chang et al., 2012; Glista et al., 2012; Baydan et al., 2019). Overall, their findings showed that aided behavioral thresholds were strongly correlated with CAEP responses to short non-speech as well as speech sounds. However, we believe that the use of continuous speech is more ecologically relevant compared to speech sounds and could potentially provide more information concerning the functionality of the hearing aid in daily life.

## 5 CONCLUSION

To summarize, the results of the present study showed that neural envelope tracking increases with increasing SNR. However, this increase is not strictly monotic as delta band tracking converges between 0 and 4 dB SNR, similarly to the behavioral outcomes. Moreover, these results were confirmed by the TRF analysis, suggesting that neural tracking correlates with speech intelligibility rather than stimulus SNR. In contrast, the ability of children’s brain to track the speech envelope in the theta band was drastically reduced and more easily corrupted by increasing noise in comparison to the delta band. Lastly, we found that neural envelope tracking in the delta band was directly associated with behavioral measures of speech intelligibility. Altogether, our findings provide a unique basis for a behavior-free evaluation of speech intelligibility in difficult-to-test populations, such as young children.

## AUTHOR STATEMENT

**Tilde Van Hirtum**: Formal analysis, investigation, data curation, writing-original draft, visualization and project administration. **Ben Somers**: Investigation, writing-review and editing, software. **Eline Verschueren**: Investigation, writing-review and editing and project administration. **Benjamin Dieudonné**: Writing-review and editing, software. **Tom Francart**: Conceptualization, methodology, writing-review and editing, supervision and funding acquisition.

## DECLARATION OF INTEREST

None.

## ACKNOWLEDGEMENTS

The authors are grateful to all the children and their parents who made time to participate in this study. The authors would also like to thank Mathilde Van Hecke for her help in data acquisition. Financial support was provided by the European Research Council (ERC) under the European Union’s Horizon 2020 research and innovation programme [grant number 637424, Tom Francart]; the KU Leuven research fund [grant number C3/20/045] and VLAIO innovation mandate (IM) [grant number HBC.2021.0203, Ben Somers].

## Notes

### Competing Interest Statement

The authors have declared no competing interest.

